# A CRISPR/Cas9 vector system for neutrophil-specific gene disruption in zebrafish

**DOI:** 10.1101/2020.07.27.223008

**Authors:** Yueyang Wang, Alan Y. Hsu, Eric M. Walton, Ramizah Syahirah, Tianqi Wang, Wenqing Zhou, Chang Ding, Abby Pei Lemke, David M. Tobin, Qing Deng

## Abstract

Tissue-specific knockout techniques are widely applied in biological studies to probe the tissue-specific roles of specific genes in physiology, development, and disease. CRISPR/Cas9 is a widely used technology to perform fast and efficient genome editing in vitro and in vivo. Here, we report a robust CRISPR-based gateway system for tissue-specific gene inactivation in zebrafish. A transgenic fish line expressing Cas9 under the control of a neutrophil-restricted promoter was constructed. As proof of principle, we transiently disrupted *rac2* or *cdk2* in neutrophils using plasmids driving the expression of sgRNAs from U6 promoters. Loss of the *rac2* or *cdk2* gene in neutrophils resulted in significantly decreased cell motility, which could be restored by re-expressing Rac2 or Cdk2 in neutrophils in the corresponding knockout background. The subcellular location of Rac activation and actin structure and stress in the context of neutrophil migration was determined in both the wild-type and *rac2* knockout neutrophils *in vivo*. In addition, we evaluated an alternative approach where the Cas9 protein is ubiquitously expressed while the sgRNA is processed by ribozymes and expressed in a neutrophil-restricted manner. Cell motility was also reduced upon *rac2* sgRNA expression. Together, our work provides a potent tool that can be used to advance the utility of zebrafish in identification and characterization of gene functions in neutrophils.

## Introduction

Over the past decade, zebrafish (*Danio rerio*) has become increasingly popular as a model system for various biological studies, including neutrophil biology^1^. Zebrafish-specific characteristics, including transparent embryos, a fast life cycle^2^, a highly conserved innate immune system^3^ as well as ease of genetic manipulation^4^ make the zebrafish perfect for live imaging and dissecting new mechanisms regulating neutrophil migration.

Tissue-specific knockout technology is an important approach to investigating gene function in vivo. However, most current gene inactivation approaches are not practical for tissue-specific loss-of-function studies in zebrafish model. The Cre/loxP site-specific recombination system was one of the earliest conditional gene modification systems^5^ and has been widely applied in transgenic mice^6^. The first Cre/loxP system in zebrafish was developed in 2005, by injecting *Cre* RNA into embryos of a floxed *gfp* transgenic line^7^. Since then, the Cre/loxP system was adapted to the zebrafish model for tissue-specific gene modification^8–10^. However, the generation of floxed alleles is complex and time-consuming, and the available cell-specific or tissue-specific Cre driver lines are limited, which restricted the utility of this technology in zebrafish^11^. Other gene-silencing approaches such as RNA interference (RNAi) is successful in some cases^12,13^, but are not generally effective in the zebrafish model^14^. Many studies reported high levels of “off-target” effects after injecting dsRNA into zebrafish embryos^15,16^. Small interfering RNAs (siRNAs) driven by U6 promoters were not functional in zebrafish^17^. Therefore, RNAi has not been widely used for studying gene function in zebrafish.

In the past few years, CRISPR/Cas9-based technology has been successfully used in zebrafish to edit the genome efficiently^18^. The Zon group designed a CRISPR-based vector system, termed miniCooper, that enabled tissue-specific gene inactivation in zebrafish^19^. Our group has adapted this system and applied it to neutrophils in zebrafish^20^. Although we achieved high efficiency of gene disruption, a major limitation is observed: knockout efficiency drops significantly when the knockout line is crossed with other lines that use the neutrophil-specific promoter. This created two problems: 1. without a sgRNA-resistant rescue construct, the specificity of the sgRNAs and the related phenotype cannot be concluded. 2. It is difficult to incorporate biosensors into the knockout lines, limiting the live imaging approaches in the context of the cell-specific knockout. We speculate that the presence of another construct driven by the same neutrophil-specific promoter in the genome competes with the transcriptional factors for Cas9 expression and reduces Cas9 protein to a level that is not sufficient for efficient knockout. In addition to using U6 promoters to drive sgRNA expression, multiple sgRNAs expression under one promoter has been achieved by utilizing the endogenous tRNA processing machinery^21^. This approach, in theory, can also be used to restrict the expression of Cas9 and sgRNAs and achieve tissue specific disruption.

Here, we report a robust CRISPR/Cas9 system for neutrophil-specific knockout in zebrafish. We successfully disrupted different genes in neutrophils and applied biosensor-based imaging in the knockout background. To test the efficacy of our system, we inactivated the *rac2* gene. Rac2 is essential for actin polymerization, migration and signaling. Loss of Rac2 activity leads to defects in neutrophil motility and chemotaxis in zebrafish^22, 23, 24^. However, the subcellular localization of Rac activation during neutrophil migration *in vivo* remains unclear. Our observation of the subcellular localization of Rac activation and related actin structure in zebrafish neutrophils sheds light on the dynamic role of Rac2 in regulating neutrophil migration.

## Results

### A Gateway cloning system for tissue-specific knockout

We optimized the Gateway system using untagged Cas9. Our strategy was to express Cas9 specifically in neutrophils and the sgRNA ubiquitously to achieve neutrophil-specific knockout. The first construct of the system, cell-specific Cas9 expression, was generated from recombination of four Gateway plasmids, containing, respectively, the neutrophil-specific promoter, Cas9 with nuclear localization sequences, SV40 polyA, and a Tol2 backbone with GFP reporter gene driven by the α*-crystallin* (*cry*) promoter. GFP positive lenses enable selection of zebrafish with stable genomic integration of this construct (Fig. 1A). The transgenic line *Tg*(*lyzC:cas9, cry:GFP*)^*pu26*^ was generated using this plasmid. The second construct of the system introduces ubiquitous sgRNA expression. This plasmid has a GFP reporter gene controlled by *lyzC* promoter, and zebrafish RNA polymerase (RNAP) III-dependent U6 promoters (U6a and U6c), each driving one gene-specific sgRNA (Fig. 1B). The successful incorporation of sgRNA sequences into the zebrafish genome can be visualized by GFP expression in neutrophils.

**Figure 1.**
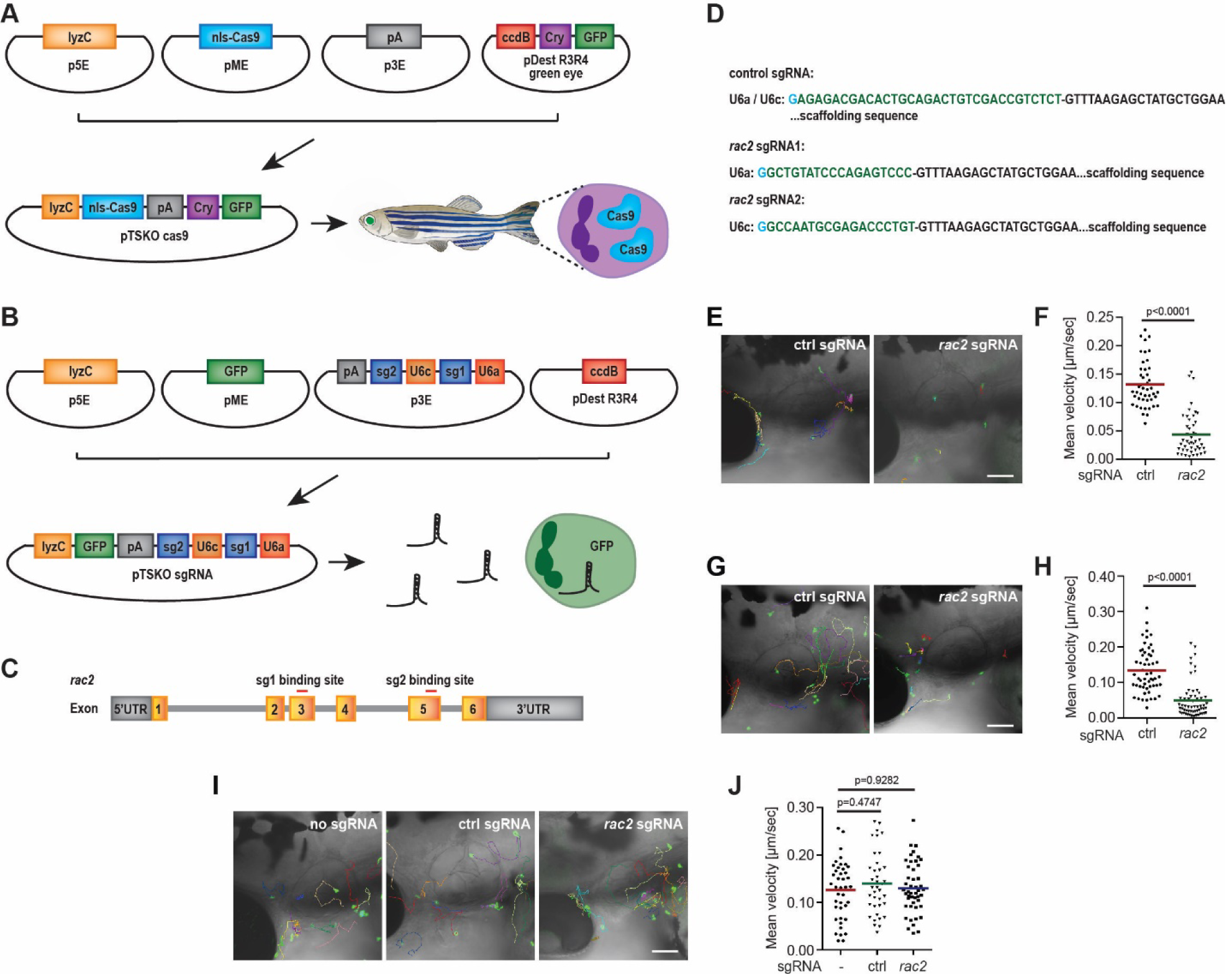
Establishment and characterization of the neutrophil-specific knockout system. (A) Schematics of the gateway vectors to clone constructs for the zebrafish *Tg*(*lyzC:Cas9, Cry:GFP*) ^*pu26*^ line with neutrophil-specific Cas9 expression and green lens. (B) Schematics of the gateway constructs for ubiquitous expression of sgRNAs and neutrophil-specific expression of GFP or other reporters. (C) Schematic of the gene structure of zebrafish *rac2* gene. The sgRNA1 and sgRNA2 target exon 3 and exon 5 in the forward strand, respectively. (D) Sequences of control and *rac2* sgRNAs. (E, F) Representative tracks (E) and quantification (F) of neutrophil motility in the head mesenchyme of 3 dpf *Tg*(*lyzC:Cas9, Cry:GFP*) ^*pu26*^ larvae injected with plasmids carrying control (ctrl) or *rac2* sgRNAs. n=45 for control, and n=46 for *rac2* transient knockouts from four different larvae. (G, H) Stable lines were generated by crossing *Tg*(*lyzC:Cas9, Cry:GFP*) ^*pu26*^ with *Tg(u6a/c: ctrl sgRNA, lyzC:GFP*)^*pu27*^ or *Tg(u6a/c: rac2 sgRNA, lyzC:GFP*) ^*pu28*^. Representative tracks (G) and quantification (H) of neutrophil motility in the head mesenchyme of 3 dpf larvae. n=55 for control from four different larvae and n=60 for *rac2* knockouts from five different larvae. P<0.0001, Mann–Whitney test. (I, J) Representative images (I) and quantification (J) of neutrophil motility in the head mesenchyme of 3 dpf larvae. n=41 for *Tg(lyzC:GFP)*, n=38 for *Tg(u6a/c: ctrl sgRNA, lyzC:GFP*)^*pu27*^, and n=47 for *Tg(u6a/c: rac2 sgRNA, lyzC:GFP*) ^*pu28*^ from three different larvae. P=0.9282 and P=0.4747 by one-way ANOVA Scale bars: 100 µm. See also Movie S1, S2, S3.

To test the efficiency of the gene knockout using this system, we injected the F2 embryos of the *Tg*(*lyzC:cas9, cry:GFP*) ^*pu26*^ line with the plasmids carrying *rac2* sgRNAs or *ctrl* sgRNAs for transient gene inactivation. The sequences of the sgRNAs are described in Fig. 1C, D. A longer sequence with no predicted binding sites in the zebrafish genome was used as a control sgRNA (Fig. 1D). As expected, we observed significantly decreased neutrophil motility in larvae of *Tg(lyzC:cas9, cry:GFP*) ^*pu26*^ fish transiently expressing sgRNAs targeting *rac2* (Fig. 1E, F and Movie S1), indicating that sufficient disruption of the *rac2* gene had been achieved. To test the knockout efficiency in stable lines, we generated transgenic lines of *Tg(U6a*/c: *ctrl sgRNAs, lyzC:GFP*) ^*pu27*^ or *Tg(U6a/*c: *rac2 sgRNAs, lyzC:GFP*) ^*pu28*^, crossed the F1 fish with Tg(*lyzC:cas9, cry:GFP*) ^*pu26*^ and quantified the velocity of neutrophils in the head mesenchyme of embryos at 3 dpf. A significant decrease of motility was observed in the neutrophils expressing Cas9 protein and *rac2* sgRNAs (Fig. 1G, H and Movie S2).

To ensure that the U6-induced sgRNAs alone did not influence neutrophil motilit*y*, we compared neutrophil motility in the transgenic lines of *Tg(U6a/c: ctrl sgRNAs, lyzC:GFP)* ^*pu27*^ or *Tg*(U6a/c: *rac2 sgRNAs, lyzC:GFP*) ^*pu28*^ with *Tg(lyzC:GFP)*^25^. All lines displayed similar neutrophil motility, indicating that the migration defects are dependent on the successful disruption of rac2 in neutrophils (Fig. 1I, J and Movie S3).

### Expression of sgRNAs-resistant rac2 rescued neutrophil migration defects resulted from rac2 disruption

To confirm that the observed decrease in neutrophil motility was due to the disruption of the *rac2* using the knockout system, we constructed plasmids carrying wild type (WT) or a dominant negative version, D57N (DN)^22^, of sgRNA-resistant *rac2* which also carries the *rac2* sgRNAs (Fig. 2A). As shown in Fig. 2B, C and Movie S4, restored expression of WT, but not the DN, Rac2 rescued the neutrophil motility defects resulted from neutrophil-specific *rac2* disruption.

**Figure 2.**
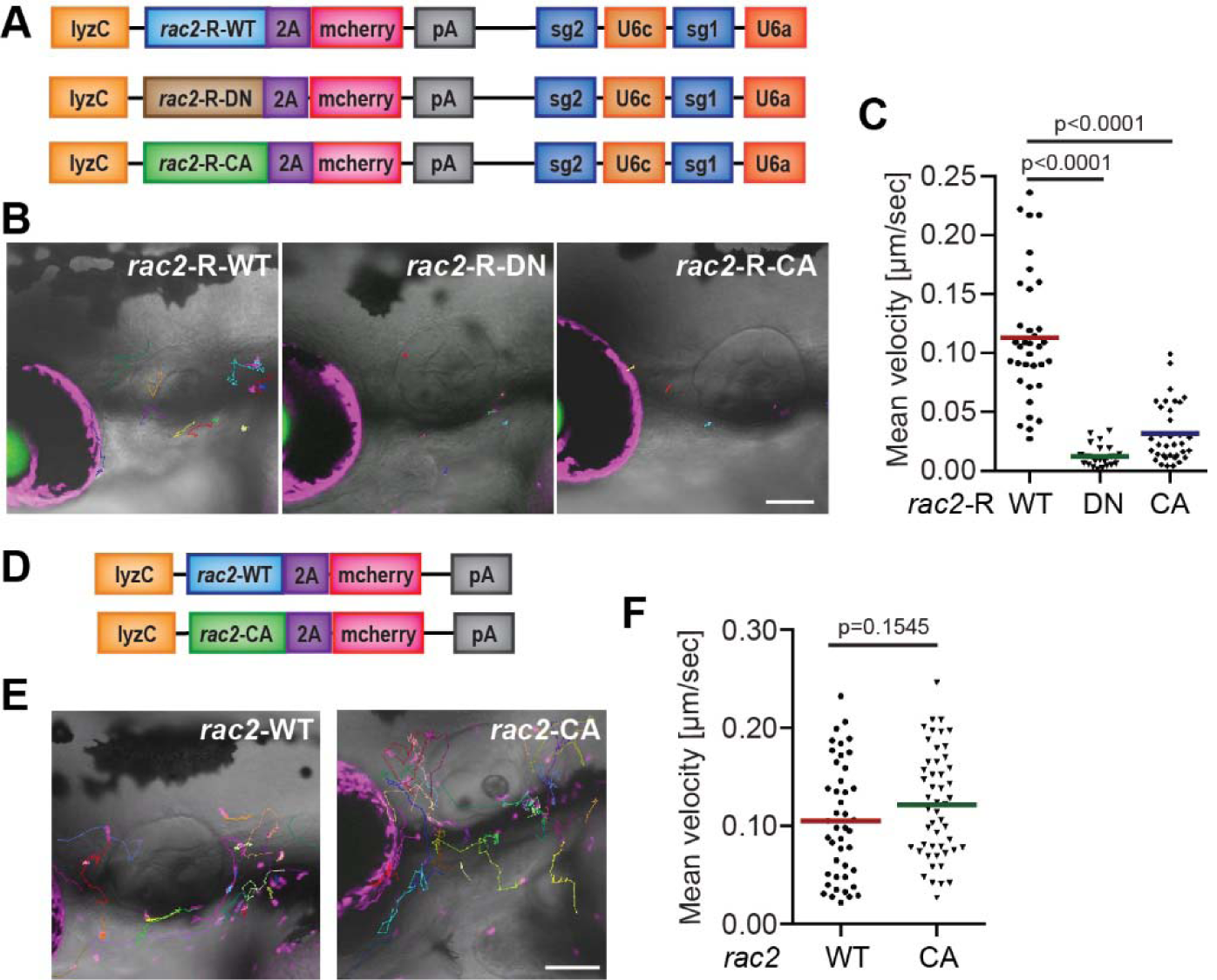
Re-expression of *rac2* rescued neutrophil motility in neutrophil-specific *rac2* knockout fish. (A) Schematics of the plasmids for neutrophil specific rescue. SgRNA-resistant *rac2* gene of wild-type (*rac2*-R-WT), dominant negative (*rac2*-R-DN), or constitutively active (*rac2*-R-CA), along with mcherry reporter gene, were cloned into the plasmid carrying *rac2* sgRNAs. Representative tracks (B) and quantification (C) of neutrophil motility in the head mesenchyme of 3 dpf *Tg*(*lyzC:Cas9, cry:GFP*) ^*pu26*^ larvae injected with plasmids containing *rac2*-R (WT), *rac2*-R-*D57N* (DN) or *rac2*-R-*Q61L* (CA). n=35 for *rac2*-R-WT, n=27 for *rac2*-R-DN and n=35 for *rac2*-R-CA from four or five different larvae respectively. P<0.0001, one-way ANOVA. (D) Schematics of the plasmids used to generate stable *Tg*(*lyzC:rac2-2A-mcherry*) ^*pu30*^or *Tg(lyzC:rac2-Q61L-2A-mcherry*) ^*pu29*^ lines. Representative tracks (E) and quantification (F) of neutrophil motility in the head mesenchyme of 3 dpf *Tg*(*lyzC:rac2-2A-mcherry*) ^*pu30*^ or *Tg(lyzC: rac2-Q61L-2A-mcherry*) ^*pu29*^ larvae. n=44 for *rac2*-WT, n=50 for *rac2*-CA from three different larvae. p=0.1545, Mann–Whitney test. Scale bars: 100 µm. See also Movie S4, S5.

A constitutively active (CA) version of RAC2, Q61L, has been reported to induce abnormal cell proliferation in human cells^26^. However, whether this mutation impacts neutrophil migration is yet to be determined. Here, we generated a transgenic zebrafish line, *Tg(lyzC:mcherry-2A-Rac2CA*) ^*pu29*^, overexpressing the Rac2 CA in neutrophils (Fig. 2D). Interestingly, in the *Tg(lyzC:mcherry-2A-Rac2CA)* ^*pu29*^ stable line, no significant change was observed regarding neutrophil motility in the F2 zebrafish larvae comparing to the that of the *Tg(lyzC:mcherry-2A-Rac2WT)* ^*pu30*^ line, which overexpresses wild-type Rac2 (Fig. 2E, F and Movie S5). On the other hand, the Rac2 CA could not rescue the motility defects resulted from neutrophil-specific *rac2* disruption (Fig. 2B, C). This indicates that the Q61L CA mutant *rac2* does not have a dominant function but only impacts neutrophil mobility in the *rac2* knockout background. Rac2 CA alone cannot coordinate neutrophil migration.

### Disruption of cdk2 using the tissue-specific knockout system also suppressed neutrophil motility

Our previous study revealed an unexpected and critical role of Cdk2 in neutrophil migration and chemotaxis^27^. To ensure that our neutrophil-specific knockout system is feasible for disrupting other genes, we injected plasmids carrying *cdk2* sgRNAs into *Tg(lyzC:cas9, cry:GFP*) ^*pu26*^ embryos. Neutrophil motility was significantly reduced by the tissue-specific *cdk2* disruption (Fig.3C, D and Movie S6), recapitulating the phenotypes observed in the stable lines overexpressing a dominant negative form of Cdk2 ^27^.

To exclude possible off-target effects for the sgRNAs targeting *cdk2* gene, we also re-expressed WT or DN versions of sgRNA-resistant *cdk2*. As shown in Fig. 3F, G and Movie S7, restored WT, but not to the DN, Cdk2 expression could partially rescue the neutrophil migration defects caused by *cdk2* gene disruption.

**Figure 3.**
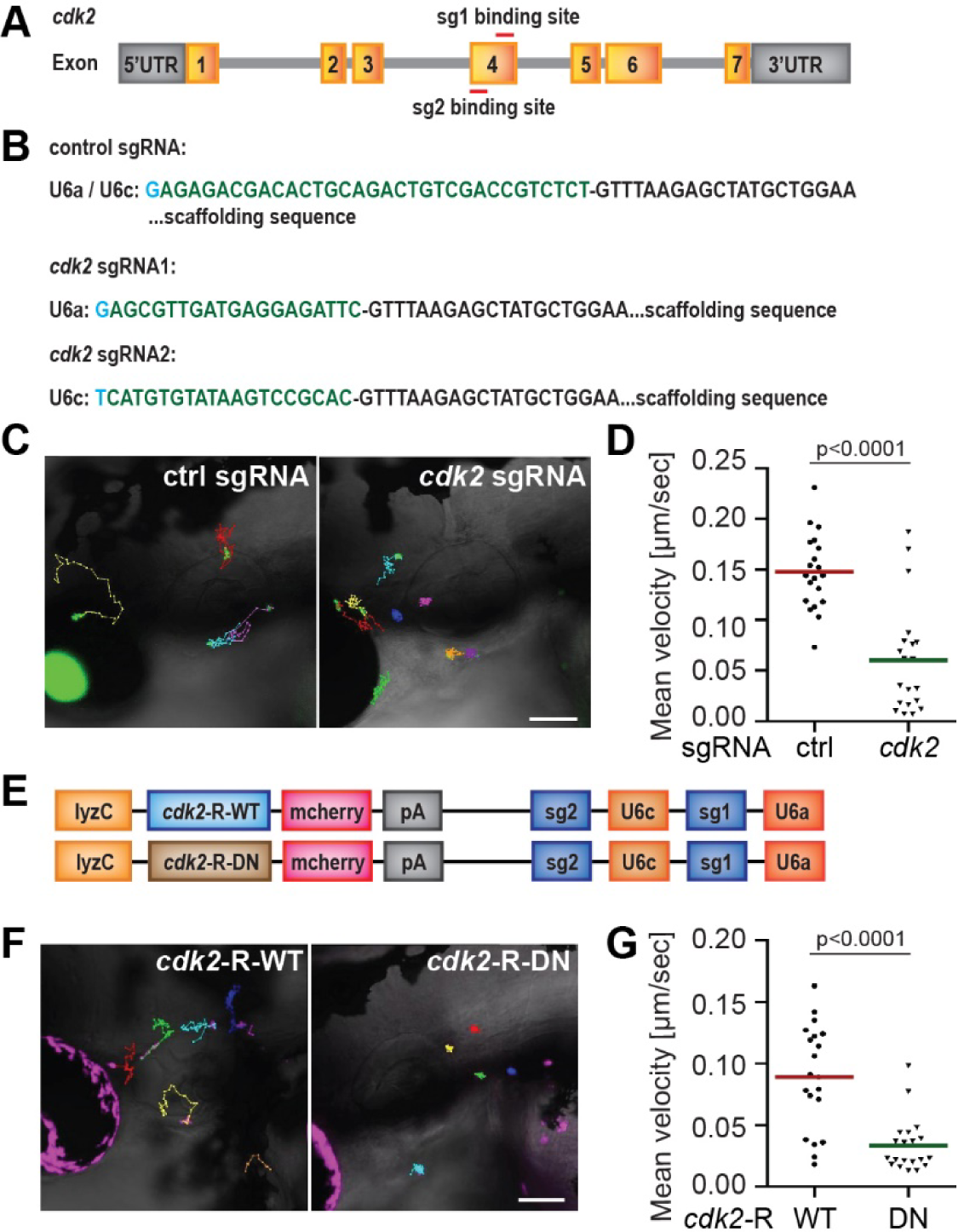
Neutrophil-specific knockout of *cdk2* reduced neutrophil motility. (A) Schematic of the gene structure of zebrafish *cdk2* gene. The two sgRNAs target sites are in exon 4. (B) Sequences of sgRNAs (control sgRNA or *cdk2* sgRNA) used in the system. Representative tracks (C) and quantification (D) of neutrophil motility in the head mesenchyme of 3dpf *Tg(lyzC:Cas9, Cry:GFP*) ^*pu26*^ larvae injected with plasmids carrying sgRNAs of control (ctrl) or *cdk2*. (E) Schematic diagrams of the plasmids used to rescue Cdk2 expression. Representative tracks (F) and quantification (G) of neutrophil motility in the head mesenchyme of 3 dpf *Tg*(*lyzC:Cas9, Cry:GFP*) ^*pu26*^ larvae injected with plasmids carrying *cdk2*-R-WT or *cdk2*-R-D145N (DN). n=20 for each group from three different larvae. ****P<0.0001, Mann– Whitney test. Scale bars: 100 µm. See also Movie S6, S7.

### Neutrophil-specific rac2 knockout disrupts Rac activation

To observe alternations of Rac activation resulted from *rac2* disruption, we used a Rac-binding domain of PAK fused with GFP (PBD-GFP)^28^ to mark the location of active Rac in neutrophils. This reporter was used in previous studies in human neutrophil-like HL-60 cells and revealed that active Rac localizes to the cell front during migration *in vitro* ^29,30,31^. Here, in zebrafish the PBD-GFP probe was enriched at both the front and rear in the migrating neutrophils expressing control sgRNAs. When the neutrophil started to migrate, Rac activity oscillates: active Rac first concentrated on the cell front and later shifted to the back. No discernible enrichment of Rac activity was detected in the *rac2* deficient neutrophils (Fig. 4B, C and Movie S8).

**Figure 4.**
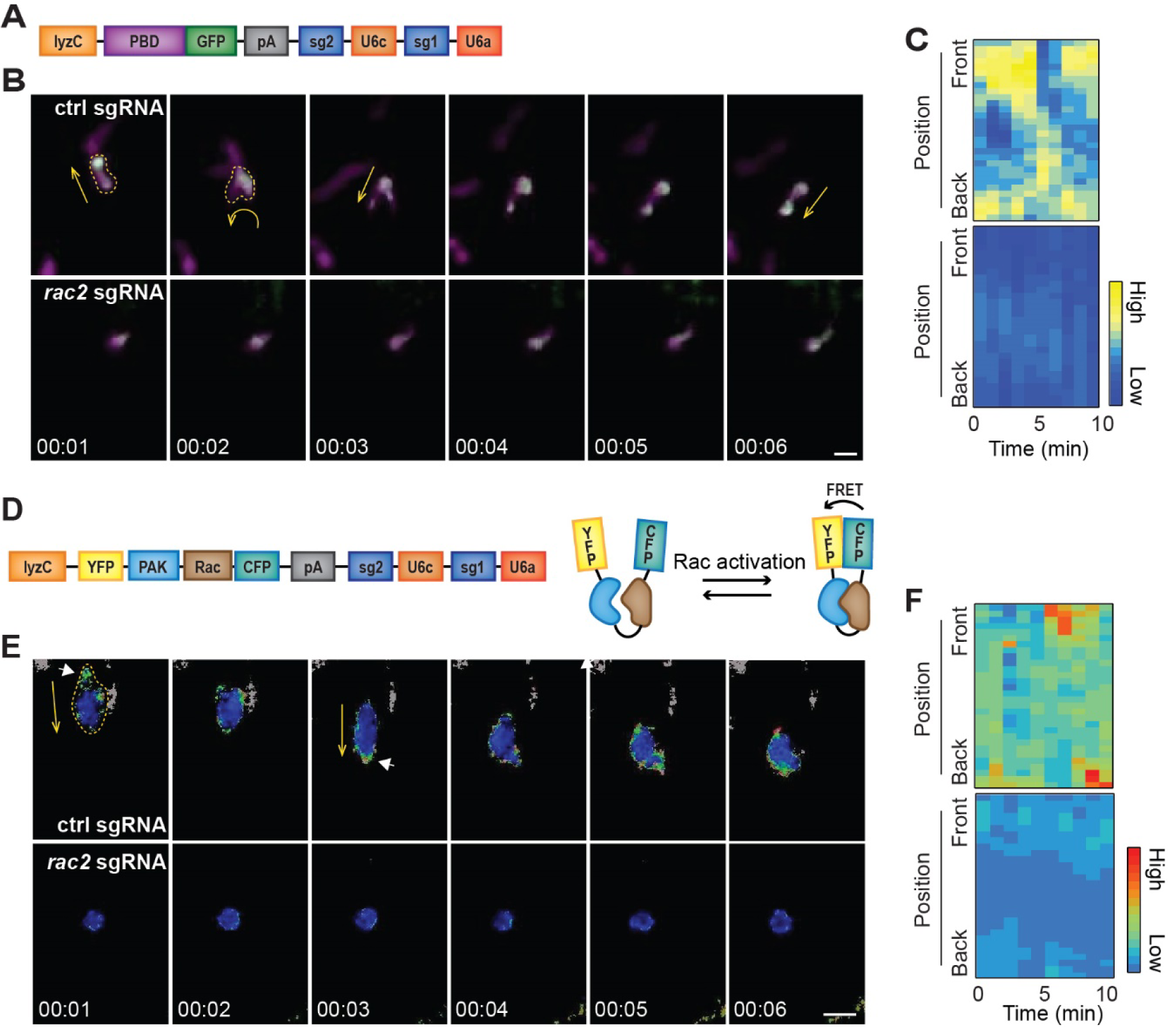
Subcellular location of Rac activation in wild-type and *rac2*-deficient neutrophils. (A) Schematic diagram of the plasmid allowing neutrophil specific PBD-GFP expression and ubiquitous sgRNA expression. (B) Simultaneous imaging of PBD-GFP and cytosolic mCherry in neutrophils expressing either ctrl sgRNA or *rac2* sgRNA. (C) Kymographs of PBD-GFP signal intensity along the axis of migration in neutrophils expressing either ctrl sgRNA (upper) or *rac2* sgRNA (lower). (D) Schematic diagram of the plasmid allowing neutrophil specific Racihu-Rac1 expression and ubiquitous sgRNA expression. (E) Representative images of Rac activity in migrating neutrophils determined by ratiometric FRET live imaging. (F) Kymographs of Racihu-Rac1 FRET intensity along the axis of migration in neutrophils expressing either ctrl sgRNA (upper) or *rac2* sgRNA (lower). Data are representative of more than 3 separate time-lapse videos. Scale bars: 10 µm. See also Movie S8, S9.

Forster resonance energy transfer (FRET)-based biosensors have been widely used to detect protein-protein interactions. To investigate the Rac activity with a second approach, the “Raichu” (Ras superfamily and interacting protein chimeric unit) Rac1-FRET probe developed by the Matsuda group^32^ was cloned into our sgRNA plasmids. Raichu is composed of CFP-RAC1-Pak-CRIB-YFP-caax. CRIB, the CDC42/Rac interactive binding motif of Pak, binds to GTP-bound Rac. When RAC is activated, the binding of RAC-GTP to Pak-CRIB will cause FRET and increase YFP/CFP fluorescence ratio (Fig. 4D). This Rac-FRET probe has been used to generate a Rac reporter mouse strain. Neutrophils were isolated from this strain and active Rac localizes at both the front and back of chemotaxing neutrophils^33^. Consistent with the observation with the PBD-GFP probe, Rac activity is higher along the cell periphery and oscillated between the front and back of migrating neutrophils expressing control sgRNAs. The *rac2* defective neutrophils lost the ability to polarize and protrude and did not display proper RAC activity (Fig. 4E, F and Movie S9).

### Neutrophil-specific rac2 knockout led to actin cytoskeletal changes

To observe the alternations in the actin cytoskeleton as a result of *rac2* disruption, we cloned genes of calponin-homology domain of utrophin (Utr-CH)-GFP^34,35,36^, which labels stable F-actin, into the sgRNA plasmids. This (Utr-Ch)-GFP has been applied and well established in leukocyte studies in zebrafish, including neutrophils^37,38^. In *Tg(lyzC:cas9, cry:GFP*)^*pu26*^ larvae transiently expressing control sgRNAs, stable F-actin was enriched at the rear of migrating neutrophils. On the contrary, *rac2* knockout neutrophils lost cell polarity and actin dynamics, as (Utr-CH)-GFP was not enriched at specific intracellular locations (Fig. 5B, C and Movie S10).

**Figure 5.**
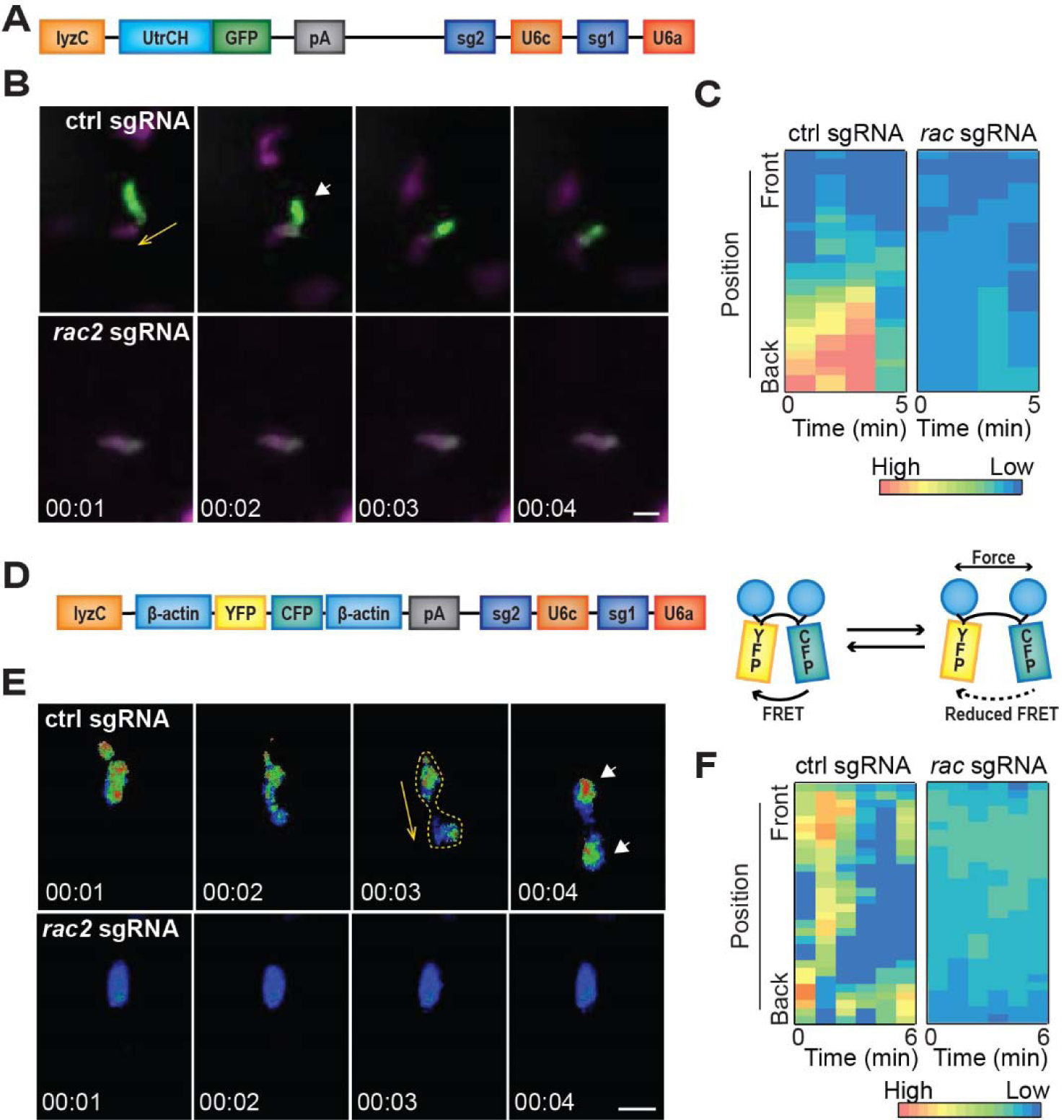
Subcellular location of stable and stresses actin in wild-type and *rac2*-deficient neutrophils. (A) Schematic diagram of the plasmid allowing neutrophil specific calponin-homology domain of utrophin (Utr-CH)-GFP expression and ubiquitous sgRNA expression. (B) Simultaneous imaging of UtrCH-GFP and cytosolic mCherry in neutrophils expressing either ctrl sgRNA or *rac2* sgRNA. (C) Kymographs of (Utr-CH)-GFP intensity along the axis of migration in neutrophils expressing either ctrl sgRNA (upper) or *rac2* sgRNA (lower). (D) Schematic diagram of the plasmid allowing neutrophil specific AcpA-FRET expression and ubiquitous sgRNA expression. (E) Ratiometric FRET live imaging of AcpA-FRET in neutrophils expressing either ctrl sgRNA or *rac2* sgRNA. (F) Kymographs of AcpA-FRET intensity along the axis of migration in neutrophils expressing either ctrl sgRNA (upper) or *rac2* sgRNA (lower). Data are representative of more than 3 separate time-lapse videos. Scale bars: 10 µm. See also Movie S10, S11.

An actin FRET probe named actin–cpstFRET–actin (AcpA) was designed by Meng group to report real-time forces within F-actin filaments^33^. The sensor consists of a FRET pair flanked with β-actin monomers on each side with protein linkers. After incorporating into F-actin filaments, the mechanical force in actin twists AcpA and decreases FRET efficiency and YFP intensity. Thus, the real-time Actin stress can be reflected with CFP/YFP ratio (Fig.5D).

We incorporated the AcpA probe into our neutrophil-specific knockout system and expressed the sensor along with control or *rac2* sgRNAs in *Tg*(*lyzC:cas9, cry:GFP*) ^*pu26*^ embryos. As shown in Fig. 5E, F and Movie S11, the actin stress FRET reporter revealed more focused actin stress at the front and rear of neutrophils in control cells during migration, while *rac2* knockout neutrophils showed decreased actin stress, suggesting that Rac2 is required for actin polymerization and force generation.

Taken together, the combination of the neutrophil-specific knockout system with various biosensors allows live imaging of the dynamic signaling events during cell migration in the knockout background.

### Ribozymes mediated gRNA generation for neutrophil-specific knockout

We also evaluated another gateway system for neutrophil-specific gene modification, in which the Cas9 protein is ubiquitously expressed while the sgRNA is processed by ribozymes and expressed in a neutrophil-restricted manner^39^. This strategy was adapted from a previous study utilizing a universal promoter and an all-in-one plasmid^40^ and we separated the Cas9 and sgRNA into two plasmids. The pME vector contains Hammerhead (HH) and hepatitis delta virus (HDV) ribozymes to cleave the RNA at the 5’ and 3’ ends respectively (Fig. 6A). The non-coding RNA, MALAT1, forming a triple helical structure at the 3′ end^41,42^, is also incorporated to stabilize the reporter gene mRNA, which is not polyadenylated. After Gateway recombination with the neutrophil-restricted promoter, a sgRNA plasmid is obtained that allows neutrophil-specific expression of the sgRNA, together with a red fluorescent protein, tdTomato.

**Figure 6.**
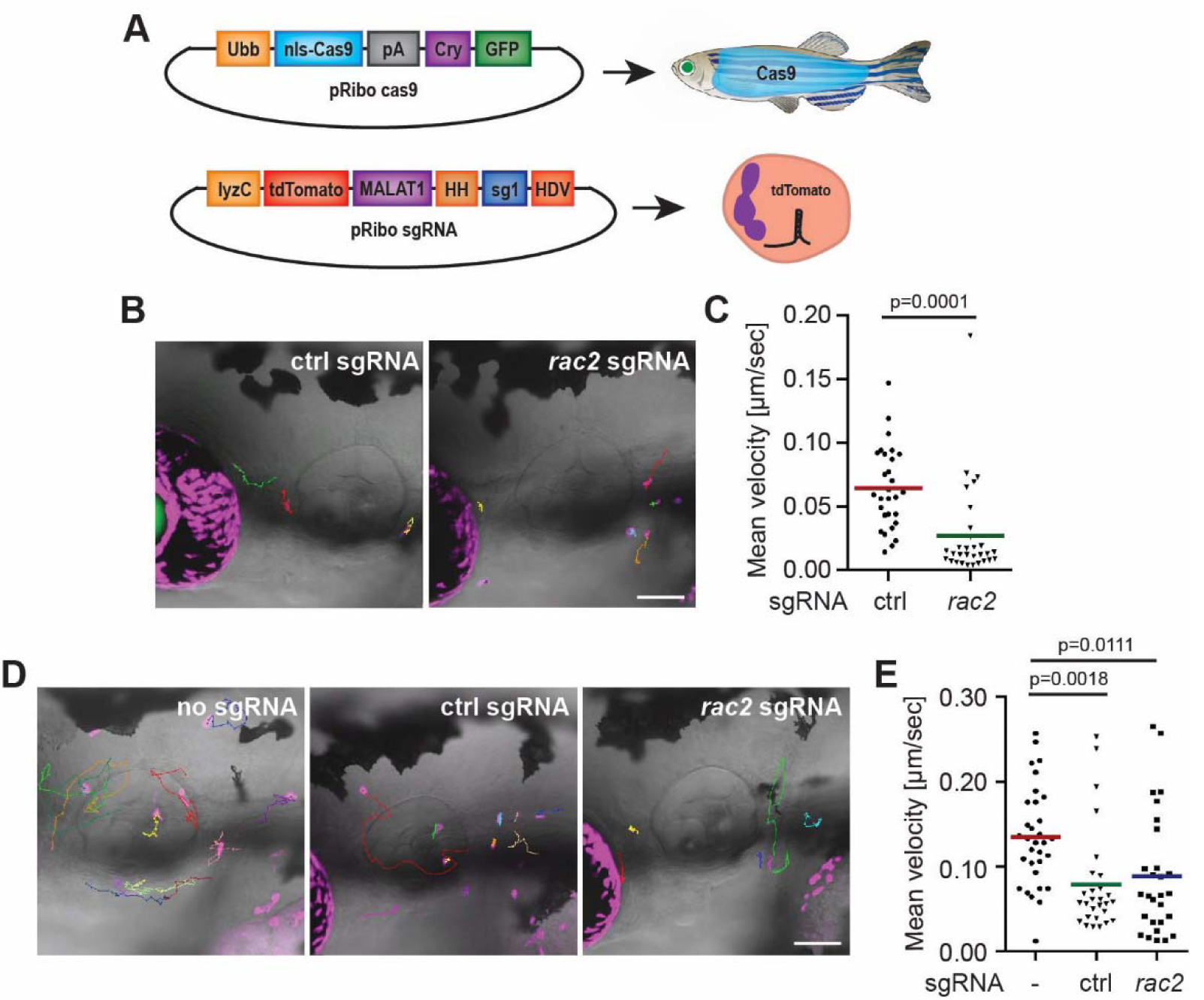
Neutrophil-specific expression of ribozyme-processed *rac*2-targeting sgRNAs reduced neutrophil motility. (A) Schematic of the design of plasmids for another neutrophil-specific knockout system. Cas9 is expressed ubiquitously whereas a sgRNA is expressed only in neutrophils. Representative tracks (B) and quantification (C) of neutrophil motility in the head mesenchyme of 3 dpf *Tg*(*ubb:Cas9, cry:GFP*)^*xt48*^ larvae injected with plasmids carrying sgRNAs of control (ctrl) or *rac2*. n=29 for control and n=30 for *rac2* transient knockouts from three different larvae. P=0.0001, Mann–Whitney test. Representative tracks (D) and quantification (E) of neutrophil motility in the head mesenchyme of 3 dpf wild-type AB zebrafish larvae injected with Tol2-lyzC-RFP plasmid or plasmids carrying sgRNAs of control (ctrl) or *rac2*. n=32 for no sgRNA control from three different larvae, n=29 for control sgRNA, and n=28 for *rac2* sgRNA from four different larvae. P=0.0018 and P=0.0111 by one-way ANOVA (E). Scale bar: 100 µm. See also Movie S12, Movie 13.

We generated a transgenic zebrafish line, *Tg(ubb:cas9, cry:GFP*)^*xt48*^ and incorporated the same control guide and the *rac2* sgRNA1 into the ribozyme-mediated knockout system. The sgRNA plasmids carrying *rac2* sgRNA or *ctrl* sgRNA were injected into the F2 embryos of *Tg(ubb:cas9, cry:GFP*) ^*xt48*^ line. As shown in Fig.6 B, C and Movie S12, significantly decreased neutrophil motility was observed in the zebrafish larvae carrying *rac2* sgRNA, indicating that sufficient gene disruption was also achieved. Notably, we observed a slight decrease in neutrophil motility when we transiently injected the control sgRNA plasmids into the wild-type background (Fig. D, E and Movie S13), which indicates potential effects the ribozyme-mediated sgRNA processing machinery may have on neutrophils.

## Discussion

Here we report a robust neutrophil-specific knockout system in zebrafish. Using a fluorescent phenotype resulting from eye-restricted GFP expression, we can easily select out the positive fish carrying Cas9 protein. Transient expression of sgRNAs and visualization of edited cells can be achieved by injecting plasmids containing sgRNAs and GFP into the Cas9-expressing fish embryos. We also demonstrated the efficiency with sgRNAs against *rac2* and *cdk2* by measuring neutrophil motility. To prove the specificity of our sgRNAs, we rescued the migration phenotype by expressing sgRNA-resistant *rac*2 or *cdk2*. The stable lines expressing Cas9 protein or *rac2* sgRNAs constructed here showed an inheritable ability to generate *rac2* knockout lines to the F2 generation. It still needs to be determined if Cas9 expression will be silenced in subsequent generations.

The optimization of the CRISPR/Cas9 vectors from our previous research lies in 1) removing the 2A-mcherry tag and 2) separating the Cas9 and sgRNA elements into two constructs. We expect that the Cas9-driver line can be maintained and crossed with different reporter lines with different sgRNAs to achieve high efficiency multiplexed knockout. With three lyzC promoter-driven constructs in one neutrophil, we were still able to observe the expected phenotypes (Figs 4B and 5B). This optimization also allows easy and flexible incorporation of other elements, such as different types of biosensors and gene specific rescue.

This system allowed novel insights into the role of the constitutively active Rac2 in neutrophil migration. We observed that the phenotypes of CA Rac2 mutant, Q61L, are different in the Rac2 wide-type and knockout background. The CA mutant can only abolish cell mobility in neutrophils without the endogenous wide-type Rac2 expression, which indicates that it cannot function dominantly in the regulation of cell migration.

It is known that the alignment of actin stress fibers in the cell body triggers cell polarization to precede cell migration. However, actin networks and the actin stress distribution in living cells, especially in a 3D tissue environment *in vivo* is not well understood. Our work provides the first observation of actin force in migrating neutrophils in vivo. We observed more focused mechanical force in actin at both the leading edge and trailing edge. The actin stress at the cell front could be attributed to the machinery driving the lamellipodium forward, which are actin networks polymerizing toward the direction of membrane protrusion and generating mechanical force to overcome membrane tension^43,44^. The actin stress at the trailing edge can be explained by the actomyosin pulling forces which facilitate the cell retraction^45^. However, how retraction of the cell rear is controlled and how the actin stress plays the role in completing the migration cycle is not well understood^46^.

We also provide the first evidence that active Rac oscillates between the front and back of migrating neutrophils *in vivo*. Previously, the characterization of Rac activity localization was limited to *in vitro* studies on two-dimensional (2D) surfaces^33^. The lamellipodium-based migration organized on monolayers on flat 2D surfaces represents only one of many mechanisms of cell migration in three-dimensional (3D) environments in tissues^47,48^. Due to the distinct modes and complexity of 3D migration, the subcellular location of Rac and its function at different sites *in vivo* was unclear. According to the currently recognized local-excitation global-inhibition model in cell migration, the two opposing processes, local-excitation and global inhibition, balance each other. Upon stimulation, the rapid excitation response overcomes the slower inhibitory signals and initiates sharp internal differences. Rac is considered as one of the front-located proteins that amplify internal asymmetries and lead to further actin polarization induced by directional sensing signals^49,50^. By using the PAK-GFP reporter for Rac, here we showed that the Rac were not only enriched at the cell membrane of the leading-edge during protrusion, but also shifted to the trailing-edge during the retraction. Fluorescence resonance energy transfer imaging of Rac activity further confirmed this subcellular localization and indicates the potential involvement of Rac in tail contraction during neutrophil migration. Our observation of subcellular location of active Rac in zebrafish neutrophils in tissue contributes to a better understanding of Rac signaling *in vivo*.

Notably, the Rac biosensors we used in theory bind to both Rac1 and Rac2 proteins^51^. Rac2 is the predominant isoform in human neutrophils^52^. In murine neutrophils, the amounts of Rac1 and Rac2 are similar^53^ and Rac1 plays a major role in mediating tail retraction^54^. In zebrafish neutrophils, Rac2 is also the dominant isoform^23,55^. It will be interesting and helpful to further look into their unique functions and signaling pathways in neutrophil migration.

Another neutrophil-specific knockout system was also assessed here. We tried applying ribozymes to generate gRNA in zebrafish neutrophils. Combined with the stable line expressing ubiquitous Cas9, this method also provides a viable approach for generating conditional mutants. For unknown reasons, fewer neutrophils are labeled using plasmid-based transient expression, and, in this case a decrease in neutrophil motility was observed with the expression of sgRNAs. An advantage of this system is that the same driver line, which drives ubiquitous Cas9 expression, can be used to knock out genes in different tissues. Further optimization, such as allowing expression of multiple sgRNAs in the system, can be attempted. Both microRNA based and tRNA based processing machinery to process multiple sgRNAs from one transcript can be incorporated^56,57,58^.

In summary, we established a robust neutrophil-specific knockout system for zebrafish. By using this system, we gained insight into the role of Rac2 in regulating the actin cytoskeleton and the subcellular location of Rac activation in zebrafish neutrophils. Our system is suitable for various genetic studies and screens, which can be achieved by injecting different sgRNAs into the Cas9-expression fish embryos. We also expect that our system can be adapted for gene function studies in other tissues by using different tissue-specific promoters.

## Materials and Methods

### Animals

The zebrafish experiment was conducted in accordance with internationally accepted standards. The Animal Care and Use Protocol was approved by the Purdue Animal Care and Use Committee (PACUC), adhering to the Guidelines for Use of Zebrafish in the National Institutes of Health (NIH) Intramural Research Program (protocol number 1401001018). To generate transgenic zebrafish lines, plasmids with the Tol2 backbone were coinjected with Tol2 transposase mRNA into embryos of the AB strain at one-cell stage as described^59^.

### Plasmids

All plasmid constructs were generated by gateway cloning using LR Clonase II Plus enzyme (Invitrogen). The p5E-lyzC entry vector was generated by replacing the ubiquitin promoter with the *lyzC* promoter in pENTR5′_ubi (Addgene #27320) as described^20^. The pME-Cas9 plasmid is from Addgene (Addgene #63154). The pME-GFP, p3E-polyA and the destination vector for sgRNA plasmid, pDestTol2pA2, are from the Tol2Kit^60^. To design single guide RNAs, CRISPRscan (http://www.crisprscan.org/) were used. Single guide RNAs with the highest score and without any off targets were selected. The destination vector pDestTol2pACryGFP (Addgene #64022) was used to generate the final Cas9 plasmids. The p3E-U6a-U6c plasmids containing *rac2* sgRNAs or control sgRNAs were same as the ones in previous work^20^, In-Fusion cloning (In-Fusion HD Cloning Plus Kit, Clontech) was used to fuse the fragments with the linearized backbone, including the wild type, DN or CA guides-resistant *rac2* (ENSDARG00000038010), or different biosensor elements. The In-Fusion primers are listed below:

pME-reverse remove GFP F:

5’-GCGGCCGCGGTGGAGCTCCAG-3’

pME-reverse remove GFP R:

5’-GGTGGCGAGTCGACCTCGAGGGG-3’

Rac-WT-guide1muta-pME F:

5’-TCTAGGCCTATGGGACACCGCAGGCCAAGAAGATTATGACAGACTGCGGCC-3’

Rac-WT-guide1muta-pME R:

5’-TCCCATAGGCCTAGATTTACCGGCTTGCTATCCACCATTACATTTGCAGAG-3’

Rac-DN-guide1muta-pME-F:

5’-TAGGCCTATGGAACACCGCAGGCCAAGAAGATTATGACAGACTGCGGCC-3’

Rac-DN-guide1muta-pME-R:

5’-TGTTCCATAGGCCTAGATTTACCGGTTTGCTATCCACCATTACATTTGC-3’

Rac-CA-guide1muta-pME-F:

5’-TCTAGGCCTATGGGACACCGCAGGCCTGGAAGATTATGACAGACTGCGG-3’

Rac-CA-guide1muta-pME-R:

5’-TCCCATAGGCCTAGATTTACCGGTTTGCTATCCACCATTACATTTGC-3’

Rac-guide2-resistant mutant F:

5’-GCAAGGACTTGCCTTAGCAAAGGAAATAGATGCAGTAAAATACCTGG -3’

Rac-guide2-resistant mutant R:

5’-AAGGCAAGTCCTTGCGGATAAGTGATCGGTGCCAGTTTC-3’

Rac-resistant into pME F:

5’-CGGGCCCCCCCTCGAGGCCACCATGGTGAGCAAGG-3’

Rac-resistant into pME R:

5’-AGCTCCACCGCGGCCGCTTAGAGCATCACGCAGCCC-3’

Rac-rescue-pME-F:

5’-GGTCGACTCGCCACCATGGTGAGCAAGGGCGAGG-3’

Rac-rescue-pME-R:

5’-CTCCACCGCGGCCGCTTAGAGCATCACGCAGCCC-3’

Rac-pME-R:

5’-AGCTCCACCGCGGCCGCTTAGAGCATCACGCAGCCC-3’

Cdk2-guide-1 F:

5’-AAGCCTCAAAACCTGCTGATCAACGCTCAGGGCGAGATC-3’

Cdk2-guide-1 R:

5’-CAGCAGGTTTTGAGGCTTAAGATCTCTGTGAAGAACCCG-3’

Cdk2-guide-1 mutant:

5’-CGGGTTCTTCACAGAGATCTTAAGCCTCAAAACCTGCTG-3’

Cdk2-guide-2 F:

5’-TACACCCACGAAGTTGTAACTTTGTGGTACAGAGCTCC-3’

Cdk2-guide-2 R:

5’-AACTTCGTGGGTGTAAGTCCGCACAGGTACACCGAACGC-3’

Cdk2-guide-2 mutant:

5’-GCGTTCGGTGTACCTGTGCGGACTTACACCCACGAAGTT-3’

ActinFRET-F-pME:

5’-GGTCGACTCGCCACCATGGATGATGATATCGCCGCGC-3’

ActinFRET-R-pME:

5’-CTCCACCGCGGCCGCCACCGCGGCCGCTTTAGAAG-3’

Utrophin-pME F:

5’-GGTCGACTCGCCACCATGGCCAAGTATGGAGAACATGAAG-3’

Utrophin-pME R:

5’-CTCCACCGCGGCCGCTTACTTGTACAGCTCGTCCATGCC-3’

PBD-F-pME:

5’-GGTCGACTCGCCACCAATACAAGCTACTTGTTCTTTTTGC-3’

PBD-R-pME:

5’-CTCCACCGCGGCCGCCTACGTAATACGACTCACTATAG-3’

RacFRET-F-pME:

5’-GGTCGACTCGCCACCATGGTGAGCAAGGGCGAGG-3’

RacFRET-R-pME:

5’-CTCCACCGCGGCCGCTTACATAATTACACACTTTGTC-3’

Ribo Ctrl F:

5’-GTATTGGTCTGCGAGAGACTGCTGATGAGTCCGTGAGGACGAAACGAGTAAGCTCG TC-3’

Ribo Ctrl R:

5’-ACTTGCTATTTCTAGCTCTAAAACGAGACGACACTGCAGACTGGACGAGCTTACTCG TT-3’

Ribo Rac guide F:

5’-GTATTGGTCTGCGAGCACTCCCTGATGAGTCCGTGAGGACGAAACGAGTAAGCTCGT C-3’

Ribo Rac guide R:

5’-ACTTGCTATTTCTAGCTCTAAAACCAGACGGTAAACACCACTCCGACGAGCTTACTC GTT-3’

### Microinjection

Microinjections were performed as described^59^. We injected 1 nL of a mixture containing 25 ng/µl plasmid and 35 ng/µl Tol2 transposase mRNA dissolved in an isotonic solution into the cytoplasm of embryos at the one-cell stage. The stable lines were generated as described^59^. At least 2 founders (F0) for each line were obtained. Experiments were performed with F2 larvae produced by F1 fish derived from multiple founders.

### Live imaging

Larvae at 3 dpf were placed on a glass-bottom dish, and imaging was performed at 28 °C. Time-lapse fluorescence images for neutrophil motility were obtained by a laser scanning confocal microscope (LSM 710, Zeiss) with a 20× objective at 1-min intervals for 30 mins. The green and red channels were acquired sequentially with 0.3 ∼ 3% power of lasers. Neutrophil migration with the expression of biosensors was captured every 1 min for 10 min. For (Utr-CH)-GFP and PBD-GFP, the green and red channels were acquired with 0.3 ∼ 5% power of the 488-nm laser and 0.5 ∼ 2% power of the 561-nm laser, respectively. The 458-nm laser channel with 75% power was used for the AcpA-FRET and Rac-FRET biosensors. The fluorescent stacks were flattened using the maximum intensity projection and overlaid with or without a single slice of the bright-field image. The velocity of neutrophils was quantified using ImageJ with MTrackJ plugin and plotted in Prism 6.0 (GraphPad). The fluorescence intensity quantification was done by an algorism written in our lab (https://github.com/tomato990/subcellular-intensity-reader). The kymograph was generated using Helm 1.0^61^.

### Statistical analysis

Statistical analysis was performed with Prism 6 (GraphPad). An unpaired two-tailed Student’s *t-* test or one-way ANOVA was used to determine the statistical significance of differences between groups. A *P*-value less than 0.05 was considered as statistically significant. Individual *P* values are indicated in the figures, with no data points excluded from statistical analysis. One representative experiment of at least 3 independent repeats is shown.

## Acknowledgements

Raichu-Rac1 is a generous gift from Dr. Miki Matsuda (Kyoto University, Japan). The work was supported by National Institutes of Health [R35GM119787 to DQ], [AI125517, AI130236, AI127115 to DT] and [P30CA023168 to Purdue Center for Cancer Research] for shared resources. AH is supported by Cagiantas Fellowship, Purdue University.

## Author contributions

YW, AH, EW, DT and DQ designed research and wrote the manuscript. YW and AH performed most experiments and analyzed the data. EW, RS, TW, WZ helped with experiments. AL and CD helped with data analysis. All authors read and approved the manuscript.

## Data availability statement

Plasmids will be available on Addgene.

## References

1. Deng, Q. & Huttenlocher, A. Leukocyte migration from a fish eye’s view. Journal of Cell Science 125, 3949–3956 (2012).

2. Driever, W., Stemple, D., Schier, A. & Solnica-Krezel, L. Zebrafish: genetic tools for studying vertebrate development. Trends in Genetics 10, 152–159 (1994).

3. Lieschke, G. J. & Trede, N. S. Fish immunology. Current Biology 19, (2009).

4. Lawson, N. D. & Wolfe, S. A. Forward and Reverse Genetic Approaches for the Analysis of Vertebrate Development in the Zebrafish. Developmental Cell 21, 48–64 (2011).

5. Hoess, R. H. & Abremski, K. Mechanism of strand cleavage and exchange in the Cre-lox site-specific recombination system. J. Mol. Biol. 181, 351–362 (1985).

6. Branda, C. S. & Dymecki, S. M. Talking about a revolution: The impact of site-specific recombinases on genetic analyses in mice. Developmental Cell 6, 7–28 (2004).

7. Pan, X., Wan, H., Chia, W., Tong, Y. & Gong, Z. Demonstration of site-directed recombination in transgenic zebrafish using the Cre/loxP system. Transgenic Res. 14, 217–223 (2005).

8. Thummel, R. et al. Cre-mediated site-specific recombination in zebrafish embryos. Dev. Dyn. 233, 1366–1377 (2005).

9. Xiong, F., Wei, Z. Q., Zhu, Z. Y. & Sun, Y. H. Targeted Expression in Zebrafish Primordial Germ Cells by Cre/loxP and Gal4/UAS Systems. Mar. Biotechnol. 15, 526–539 (2013).

10. Hans, S., Kaslin, J., Freudenreich, D. & Brand, M. Temporally-Controlled Site-Specific Recombination in Zebrafish. PLoS One 4, e4640 (2009).

11. Mosimann, C. & Zon, L. I. Advanced zebrafish transgenesis with Tol2 and application for Cre/lox recombination experiments. in Methods in Cell Biology 104, 173–194 (Academic Press Inc., 2011).

12. De Rienzo, G., Gutzman, J. H. & Sive, H. Efficient shRNA-Mediated Inhibition of Gene Expression in Zebrafish. Zebrafish 9, 97–107 (2012).

13. Dong, M. et al. Heritable and Lineage-Specific Gene Knockdown in Zebrafish Embryo. PLoS One 4, e6125 (2009).

14. Kelly, A. & Hurlstone, A. F. The use of RNAi technologies for gene knockdown in zebrafish. Brief. Funct. Genomics 10, 189–96 (2011).

15. Oates, A. C., Bruce, A. E. E. & Ho, R. K. Too much interference: Injection of double-stranded RNA has nonspecific effects in the zebrafish embryo. Dev. Biol. 224, 20–28 (2000).

16. Zhao, Z., Cao, Y., Li, M. & Meng, A. Double-stranded RNA injection produces nonspecific defects in zebrafish. Dev. Biol. 229, 215–223 (2001).

17. Wang, L. et al. U6 promoter-driven siRNA injection has nonspecific effects in zebrafish. Biochem. Biophys. Res. Commun. 391, 1363–1368 (2010).

18. Varshney, G. K. et al. High-throughput gene targeting and phenotyping in zebrafish using CRISPR/Cas9. Genome Res. 25, 1030–1042 (2015).

19. Ablain, J., Durand, E. M., Yang, S., Zhou, Y. & Zon, L. I. A CRISPR/Cas9 vector system for tissue-specific gene disruption in zebrafish. Dev. Cell 32, 756–764 (2015).

20. Zhou, W. et al. Neutrophil-specific knockout demonstrates a role for mitochondria in regulating neutrophil motility in zebrafish. DMM Disease Models and Mechanisms 11, (2018).

21. Shiraki, T. & Kawakami, K. A tRNA-based multiplex sgRNA expression system in zebrafish and its application to generation of transgenic albino fish. Sci. Rep. 8, 1–14 (2018).

22. Deng, Q., Yoo, S. K., Cavnar, P. J., Green, J. M. & Huttenlocher, A. Dual Roles for Rac2 in Neutrophil Motility and Active Retention in Zebrafish Hematopoietic Tissue. Dev. Cell 21, 735–745 (2011).

23. Rosowski, E. E., Deng, Q., Keller, N. P. & Huttenlocher, A. Rac2 Functions in Both Neutrophils and Macrophages To Mediate Motility and Host Defense in Larval Zebrafish. J. Immunol. 197, 4780–4790 (2016).

24. Hsu, A. Y. et al. Inducible overexpression of zebrafish microRNA-722 suppresses chemotaxis of human neutrophil like cells. Mol. Immunol. 112, 206–214 (2019).

25. Hall, C., Flores, M., Storm, T., Crosier, K. & Crosier, P. The zebrafish lysozyme C promoter drives myeloid-specific expression in transgenic fish. BMC Dev. Biol. 7, 42 (2007).

26. Gu, Y. et al. Biochemical and Biological Characterization of a Human Rac2 GTPase Mutant Associated with Phagocytic Immunodeficiency. J. Biol. Chem. 276, 15929–15938 (2001).

27. Hsu, A. Y. et al. Phenotypical microRNA screen reveals a noncanonical role of CDK2 in regulating neutrophil migration. Proc. Natl. Acad. Sci. U. S. A. 116, 18561–18570 (2019).

28. Benink, H. A. & Bement, W. M. Concentric zones of active RhoA and Cdc42 around single cell wounds. J. Cell Biol. 168, 429–439 (2005).

29. Benard, V., Bohl, B. P. & Bokoch, G. M. Characterization of Rac and Cdc42 activation in chemoattractant-stimulated human neutrophils using a novel assay for active GTPases. J. Biol. Chem. 274, 13198–13204 (1999).

30. Srinivasan, S. et al. Rac and Cdc42 play distinct roles in regulating PI(3,4,5)P3 and polarity during neutrophil chemotaxis. J. Cell Biol. 160, 375–385 (2003).

31. Peng, G. E., Wilson, S. R. & Weiner, O. D. A pharmacological cocktail for arresting actin dynamics in living cells. Mol. Biol. Cell 22, 3986–3994 (2011).

32. Itoh, R. E. et al. Activation of Rac and Cdc42 Video Imaged by Fluorescent Resonance Energy Transfer-Based Single-Molecule Probes in the Membrane of Living Cells. Mol. Cell. Biol. 22, 6582–6591 (2002).

33. Johnsson, A. K. E. et al. The Rac-FRET Mouse Reveals Tight Spatiotemporal Control of Rac Activity in Primary Cells and Tissues. Cell Rep. 6, 1153–1164 (2014).

34. Burkel, B. M., Von Dassow, G. & Bement, W. M. Versatile fluorescent probes for actin filaments based on the actin-binding domain of utrophin. Cell Motil. Cytoskeleton 64, 822–832 (2007).

35. Barros-Becker, F., Lam, P. Y., Fisher, R. & Huttenlocher, A. Live imaging reveals distinct modes of neutrophil and macrophage migration within interstitial tissues. J. Cell Sci. 130, 3801–3808 (2017).

36. Lam, P. Y., Fischer, R. S., Shin, W. D., Waterman, C. M. & Huttenlocher, A. Spinning disk confocal imaging of neutrophil migration in Zebrafish. Methods Mol. Biol. 1124, 219–233 (2014).

37. Barros-Becker, F., Lam, P.-Y., Fisher, R. & Huttenlocher, A. Live imaging reveals distinct modes of neutrophil and macrophage migration within interstitial tissues. (2017). doi:10.1242/jcs.206128

38. Lam, P. Y., Fischer, R. S., Shin, W. D., Waterman, C. M. & Huttenlocher, A. Spinning disk confocal imaging of neutrophil migration in Zebrafish. Methods Mol. Biol. 1124, 219–233 (2014).

39. Walton, E. M. Cell Wall Lipids Promoting Host Angiogenesis During Mycobacterial Infection. (Duke University, 2018).

40. Lee, R. T. H., Ng, A. S. M. & Ingham, P. W. Ribozyme mediated gRNA Generation for in vitro and in vivo CRISPR/Cas9 mutagenesis. PLoS One 11, (2016).

41. Wilusz, J. E., Freier, S. M. & Spector, D. L. 3′ End Processing of a Long Nuclear-Retained Noncoding RNA Yields a tRNA-like Cytoplasmic RNA. Cell 135, 919–932 (2008).

42. Wilusz, J. E. et al. A triple helix stabilizes the 3′ ends of long noncoding RNAs that lack poly(A) tails. Genes Dev. 26, 2392–2407 (2012).

43. Schaks, M., Giannone, G. & Rottner, K. Actin dynamics in cell migration. Essays Biochem. 63, 483–495 (2019).

44. Membrane Tension and Cytoskeleton Organization in Cell Motility - PubMed. Available at: https://pubmed.ncbi.nlm.nih.gov/26061624/. (Accessed: 11th June 2020)

45. Wu, J. et al. Actomyosin pulls to advance the nucleus in a migrating tissue cell. Biophys. J. 106, 7–15 (2014).

46. Hetmanski, J. H. R. et al. Membrane Tension Orchestrates Rear Retraction in Matrix-Directed Cell Migration. Dev. Cell 51, 460-475.e10 (2019).

47. Petrie, R. J. & Yamada, K. M. Multiple mechanisms of 3D migration: The origins of plasticity. Current Opinion in Cell Biology 42, 7–12 (2016).

48. Petrie, R. J. & Yamada, K. M. At the leading edge of three-dimensional cell migration. Journal of Cell Science 125, 5917–5926 (2012).

49. Franca-Koh, J. & Devreotes, P. N. Moving Forward: Mechanisms of Chemoattractant Gradient Sensing. Physiology 19, 300–308 (2004).

50. Kutscher, B., Devreotes, P. & Iglesias, P. A. Local excitation, global inhibition mechanism for gradient sensing: an interactive applet. Sci. STKE 2004, pl3–pl3 (2004).

51. Sells, M. A. et al. Human p21-activated kinase (Pak1) regulates actin organization in mammalian cells. Curr. Biol. 7, 202–210 (1997).

52. Heyworth, P. G., Bohl, B. P., Bokoch, G. M. & Curnutte, J. T. Rac translocates independently of the neutrophil NADPH oxidase components p47(phox) and p67(phox). Evidence for its interaction with flavocytochrome b558. J. Biol. Chem. 269, 30749–30752 (1994).

53. Li, S. et al. Chemoattractant-Stimulated Rac Activation in Wild-Type and Rac2-Deficient Murine Neutrophils: Preferential Activation of Rac2 and Rac2 Gene Dosage Effect on Neutrophil Functions. J. Immunol. 169, 5043–5051 (2002).

54. Filippi, M. D., Szczur, K., Harris, C. E. & Berclaz, P. Y. Rho GTPase Rac1 is critical for neutrophil migration into the lung. Blood 109, 1257–1264 (2007).

55. Tell, R. M., Kimura, K. & Palić, D. Rac2 expression and its role in neutrophil functions of zebrafish (Danio rerio). Fish Shellfish Immunol. 33, 1086–1094 (2012).

56. Port, F. & Bullock, S. L. Augmenting CRISPR applications in Drosophila with tRNA-flanked sgRNAs. Nat. Methods 13, 852–854 (2016).

57. Xie, K., Minkenberg, B., Yang, Y. & Doudna, J. A. Boosting CRISPR/Cas9 multiplex editing capability with the endogenous tRNA-processing system. doi:10.1073/pnas.1420294112

58. Wang, J. et al. Generation of cell-type-specific gene mutations by expressing the sgRNA of the CRISPR system from the RNA polymerase II promoters. Protein Cell 6, 689–692 (2015).

59. Deng, Q., Yoo, S. K., Cavnar, P. J., Green, J. M. & Huttenlocher, A. Dual Roles for Rac2 in Neutrophil Motility and Active Retention in Zebrafish Hematopoietic Tissue. Dev. Cell 21, 735–745 (2011).

60. Kwan, K. M. et al. The Tol2kit: A multisite gateway-based construction Kit for Tol2 transposon transgenesis constructs. Dev. Dyn. 236, 3088–3099 (2007).

61. Deng, W., Wang, Y., Liu, Z., Cheng, H. & Xue, Y. HemI: A Toolkit for Illustrating Heatmaps. PLoS One 9, e111988 (2014).

