## Supplementary material for "A CRISPR/Cas9 vector system for neutrophil-specific gene disruption in zebrafish": Movie legend

**Movie 1. Tracked movies of migrating neutrophils in the head mesenchyme transiently expressing control or *rac2* sgRNAs.** The video shows the motility of neutrophils in 3 dpf *Tg(lyzC:Cas9, cry:GFP*) *^pu26^* zebrafish larvae injected with plasmids carrying control or *rac2* sgRNAs. Videos were recorded for 30 min with 1min interval. Representative videos from n = 3 independent experiments with 4 fish each group are shown. Scale bar: 100 μm.

**Movie 2. Tracked movies of neutrophil motility in the head mesenchyme of the control and *rac2* knockout stable lines**. We generated stable lines by crossing *Tg*(*lyzC:Cas9, Cry:GFP*) *^pu26^* with *Tg(u6a/c: ctrl sgRNA, lyzC:GFP*)*^pu27^* or *Tg(u6a/c: rac2 sgRNA, lyzC:GFP*) *^pu28^*. The video shows the motility of neutrophils in 3 dpf zebrafish offspring larvae. Videos were recorded for 30 min with 1 min interval. Representative videos from n = 3 independent experiments with 3 fish each group are shown. Scale bar: 100 μm.

**Movie 3. Tracked movies of neutrophil motility in the head mesenchyme of the no-guide control, control and *rac2* sgRNAs stable lines**. The video shows the motility of neutrophils in 3 dpf zebrafish larvae of *Tg(lyzC:GFP*), *Tg(u6a/c: ctrl sgRNA, lyzC:GFP*)*^pu27^* or *Tg(u6a/c: rac2 sgRNA, lyzC:GFP*) *^pu28^*. Videos were recorded for 30 min with 1 min interval. Representative videos from n = 3 independent experiments with 3 fish each group are shown. Scale bar: 100 μm.

**Movie 4. Tracked movies of neutrophil motility in the head mesenchyme transiently expressing *rac2*-R-WT, *rac2*-R-DN, or *rac2*-R-CA.** The video shows the motility of neutrophils in 3 dpf *Tg*(*lyzC:Cas9, cry:GFP*) *^pu26^* zebrafish larvae injected with plasmids carrying *rac2* sgRNAs along with *rac2*-R-WT, *rac2*-R-DN, or *rac2*-R-CA. Videos were recorded for 30 min with 1 min interval. Representative videos from n = 3 independent experiments with 3 fish each group are shown. Scale bar: 100 μm.

**Movie 5. Migrating neutrophils in the head mesenchyme of the *rac2*-CA and the *rac2*-CA lines.** The video shows the motility of neutrophils in 3 dpf *Tg*(*lyzC:rac2-WT-2a-mcherry*) *^pu30^* or *Tg(lyzC:rac2-CA-2a-mcherry*) *^pu29^* zebrafish larvae. Videos were recorded for 30 min with 1 min interval. Representative videos from n = 3 independent experiments with 3 fish each group are shown. Scale bar: 100 μm.

**Movie 6. Tracked movies of migrating neutrophils in the head mesenchyme transiently expressing control or *cdk2* sgRNAs.** The video shows the motility of neutrophils in 3 dpf Tg(*lyzC:Cas9, cry:GFP*) *^pu26^* zebrafish larvae injected with plasmids carrying control or *cdk2* sgRNAs. Videos were recorded for 30 min with 1 min interval. Representative videos from n = 3 independent experiments with 3 fish each group are shown. Scale bar: 100 μm.

**Movie 7. Transient expression of *cdk2*-R-WT, not *cdk2*-R-DN, restored cell motility in *cdk2*-deficient neutrophils.** The video shows the motility of neutrophils in 3 dpf *Tg*(*lyzC:Cas9, cry:GFP*) *^pu26^* zebrafish larvae injected with plasmids containing *cdk2* sgRNAs along with *cdk2*-R-WT or *cdk2*-R-DN. Videos were recorded for 30 min with 1 min interval. Representative videos from n = 3 independent experiments with 3 fish each group are shown. Scale bar: 100 μm.

**Movie 8. Neutrophil-specific *rac2* knockout lead to deficiency in the front-to-rear localization of Rac in neutrophils**. The video shows the subcellular localization of PBD-GFP, which marks the location of Rac in neutrophils of 3 dpf *Tg*(*lyzC:Cas9, cry:GFP*) *^pu26^* zebrafish larvae injected with plasmids containing control or *rac2* sgRNAs. Cytoplasm is labeled with mCherry. Scale bar: 20 μm.

**Movie 9. Neutrophil-specific *rac2* knockout abolished the oscillation between the front and rear of active Rac in neutrophils**. The video shows the subcellular localization of Rac-FRET, of which the YFP/CFP fluorescence ratio indicates the location of active Rac in neutrophils of 3 dpf *Tg(lyzC:Cas9, cry:GFP*) *^pu26^* zebrafish larvae injected with plasmids containing control or *rac2* sgRNAs. Scale bar: 10 μm.

**Movie 10. Neutrophil-specific *rac2* knockout induced stable F-actin changes in neutrophils.** The video shows neutrophils expressing GFP-UtrCH, which labels stable F-actin, along with control or *rac2* sgRNAs in 3 dpf *Tg(lyzC:Cas9, cry:GFP*) *^pu26^* zebrafish larvae. Cytoplasm is labeled with mCherry. Scale bar: 20 μm.

**Movie 11. Neutrophil-specific *rac2* knockout abrogated the generated actin stress at the front and the back.** The video shows neutrophils expressing AcpA-FRET along with control or *rac2* sgRNAs in 3 dpf *Tg(lyzC:Cas9, cry:GFP*) *^pu26^* zebrafish larvae. The ratiometric AcpA-FRET signals report the actin force in neutrophils. Scale bar: 100 μm.

**Movie 12. Tracked movies of migrating neutrophils in the head mesenchyme transiently expressing control or *rac2* sgRNAs.** The video shows the motility of neutrophils in 3 dpf *Tg(ubb:cas9, cry:GFP*)*^xt48^* zebrafish larvae injected with plasmids carrying Ribozyme-processing machinery along with the control or *rac2* sgRNAs. Videos were recorded for 30 min with 1 min interval. Representative videos from n = 3 independent experiments with 3 fish each group are shown. Scale bar: 100 μm.

**Movie 13. Tracked movies of migrating neutrophils in the head mesenchyme of zebrafish transiently expressing the neutrophil specific RFP with or without control or *rac2* sgRNAs**. The video shows the motility of neutrophils in 3 dpf wide-type AB zebrafish larvae transiently expressing RFP with or without control sgRNA or *rac2* sgRNA in neutrophils. Videos were recorded for 30 min with 1 min interval. Representative videos from n = 3 independent experiments with 4 fish each group are shown. Scale bar: 100 μm.
